# Markov modeling reveals novel intracellular modulation of the human TREK-2 selectivity filter

**DOI:** 10.1101/092155

**Authors:** Matthew P. Harrigan, Keri A. McKiernan, Veerabahu Shanmugasundaram, Rajiah Aldrin Denny, Vijay S. Pande

**Author notes:** Denotes equal contribution (Matthew P. Harrigan, Keri A. McKiernan).

## Abstract

Two-pore domain potassium (K2P) channel ion conductance is regulated by diverse stimuli that directly or indirectly gate the channel selectivity filter (SF). Recent crystal structures for the TREK-2 member of the K2P family reveal distinct “up” and “down” states assumed during activation via mechanical stretch. We performed 195 μs of all-atom, unbiased molecular dynamics simulations of the TREK-2 channel to probe how membrane stretch regulates the SF gate. Markov modeling reveals a novel “pinched” SF configuration that stretch activation rapidly destabilizes. Free-energy barrier heights calculated for critical steps in the conduction pathway indicate that this pinched state impairs ion conduction. Our simulations predict that this low-conductance state is accessed exclusively in the compressed, “down” conformation in which the intracellular helix arrangement allosterically pinches the SF. By explicitly relating structure to function, we contribute a critical piece of understanding to the evolving K2P puzzle.

---

TREK-2 is a member of the human K2P family of tandem-pore potassium channels. This protein is responsible for leak currents in nearly all cells. Its dysregulation has been linked to pain and depression, and it can be functionally regulated by mechanical stretch, heat, fatty acids, pH, secondary messengers of signaling proteins, and several drugs [1–3]. Recent evidence suggests that the regulatory processes acting on this channel modulate conductance through structurally distinct mechanisms [4]. However, the specific details of the conformational changes involved in these mechanisms remain unclear.

Potassium channel ion conductance is mediated by both an intracellular and interior gating process. The former involves large-scale rearrangements of the intracellular domains of the channel’s transmembrane helices to sterically block passage of ions. Interior gating involves smaller conformational changes occurring directly in the channel’s selectivity filter (SF), so named for its role in conferring potassium selectivity to the channel. The SF is formed by the p-loop domains connecting the protein subunits. These p-loops are arranged parallel to one another and are radially symmetric about the ion conduction pathway such that the backbone carbonyls point inward to form 4 adjacent binding sites (Fig. 1). Interior gating alters the stability of these binding sites.

**FIG. 1.**
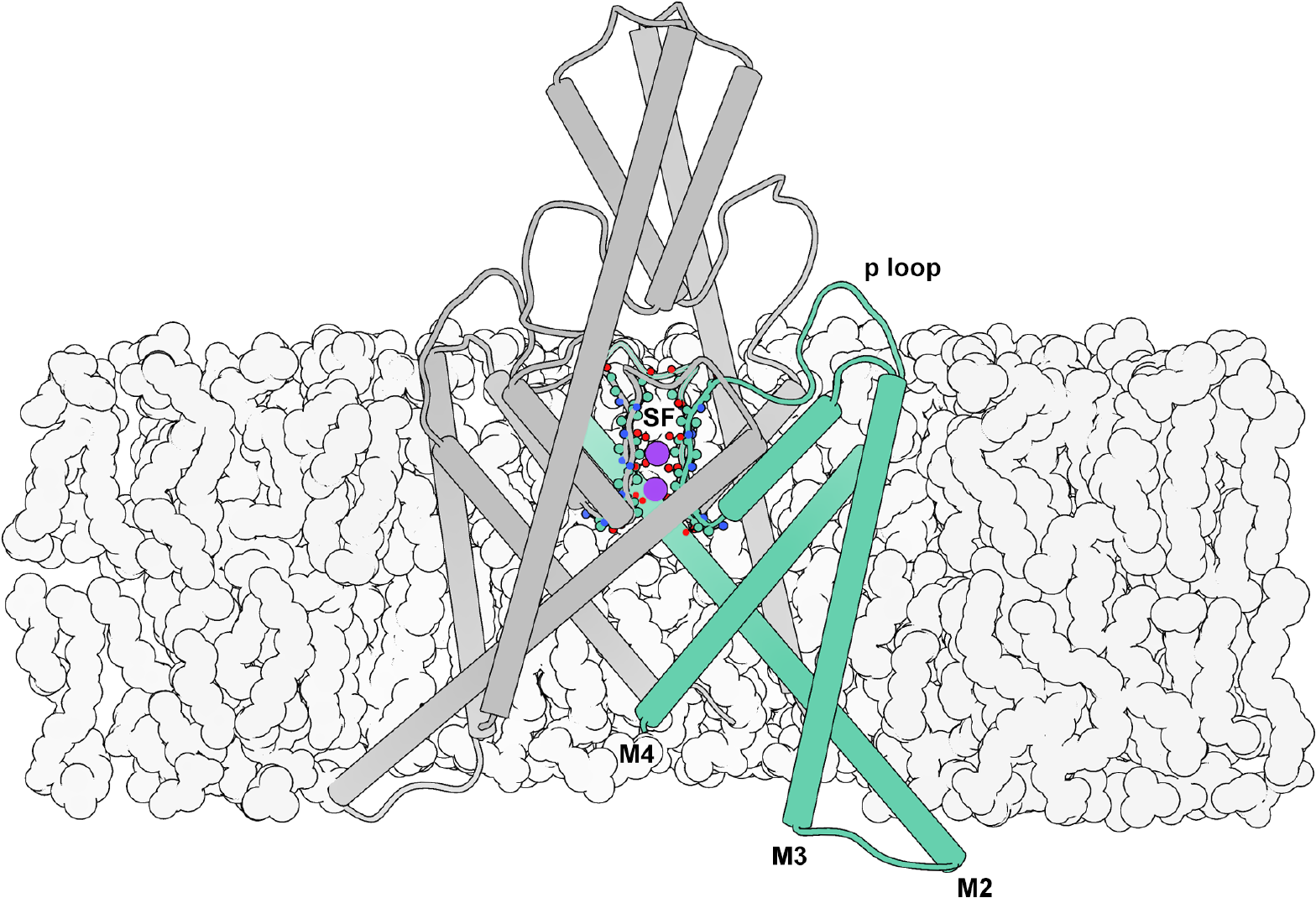
Illustration of the TREK-2 system. The selectivity filter (SF) is located at the center of the extracellular region of the membrane embedded pore, and is formed by the backbone of the four p loops. Here it is shown with occupancy corresponding to the simulated resting state. Intracellular gating involves movements of the M2, M3, and M4 helices, while interior gating involves conformational changes of the SF.

In contrast to other types of potassium channels, switching of the TREK-2 intracellular gate is not thought to “close” the channel directly. Rather, the helical conformational changes confer information to the SF, which acts as the primary gate [5–7]. Recent crystal structures of TREK-2 provide structural insight into major conformational states involved in mechanical (stretch) activation by resolving distinct “up” (stretched) and “down” (compressed) conformations [8]. The number of bound ions observed for each state suggests differing levels of conductance, rather than closure of the channel. To understand how the channel switches between these conductance levels, we apply dynamical techniques to complement static crystal structures.

Molecular dynamics (MD) is a method for simulating biological systems with atom-level resolution. By using commodity GPUs [9] paired with a massively distributed architecture [10], simulations can reach biologically-relevant timescales. Advances in computing result in large quantities of high-dimensional time-series data. To draw interpretable conclusions from this information, further analysis is required. Markov state models (MSMs) have gained favor for their robust, statistical analysis of biophysical systems [11–14]. Potential of mean force (PMF) computations can complement MSM analysis by offering quantification of free-energy profiles along paths or coordinates of interest [15].

In this paper we investigate how the structural conformational changes involved in stretch activation influence the thermodynamics and kinetics of ion permeation. We sample along this pathway through the unbiased MD simulation of the crystal structures solved by Dong *et al*. [8]. We then examine the dynamical behavior of these simulations using Markov modeling. Specifically, in Section I A, we apply MSM analysis of a large MD dataset to survey equilibrium conformational dynamics. We detect a novel “pinched” configuration of the SF favored during compression. In Section IB, we survey SF equilibrium ion dynamics, and contrast structural macrostates’ unique SF ion occupancy preferences. In Section IC, we compute PMF curves to quantitatively analyze the energetics involved in SF ion occupancy state transitions. These energetics show a correlation with both the whole-molecule and SF conformational states. Notably, the states favored under membrane compression have the most disfavored outward ion rectification. Our results predict a specific structural motif that we implicate in the coupling of intracellular and interior gates. We propose the novel conformation as a basis for future study.

## I. RESULTS

### A. Dynamics of conformational change

X-ray crystal structures offer an unrivaled atom-level view into static structures of proteins. The published crystal structures for TREK-2 identify two distinct stable conformations for the channel, presumed to be the result of membrane stretch and compression. The stretched, “up” state’s transmembrane (M) helices are displaced upwards and outwards compared to the compressed, “down” state. The conformational differences are large compared to the size of the protein. In this paper, we refer to the states distinguished by large-scale motions as “macrostates”. These static structures lack dynamical information, and the relative populations or free-energies of each macrostate in physiological conditions is unknown. We ran 195 μs of unbiased, explicit-solvent, explicit-membrane, isobaric molecular dynamics (MD) initialized with protein coordinates from the four crystal structures aiming to dynamically connect the experimentally observed structures and discover metastable or intermediate conformations inaccessible to crystallography.

Due to the large amount of data among many distributed trajectories, we apply state-of-the-art MSM modeling methods to glean insight from the large dataset. We use time-structure based independent component analysis (tICA) to automatically discover an “up-down” stretch activation coordinate from a large number of atom-pair distances. tICA is a machine learning technique to find the slowly-decorrelating modes in highdimensional time-series data. Biophysical processes of interest are often those that occur with the slowest timescales. By using tICA on a large set of input features, we avoid codifying our preconceived notions about the dynamics of the system into the analysis while still reducing the dimensionality of the system by projecting conformational features onto a small number (≈ 5) of kinetically-motivated independent components (tICs).

Building an MSM on these coordinates permits generation of a 500 ps representative trajectory. By inspection, we identify the primary tIC as an “up-down” measurement. The up-down trace for the MSM trajectory is shown in Fig. 2A and a movie is available (see Fig. SI and supporting files). The kinetic information in the MD dataset can be exploited to assign each observed conformation to a macrostate (Fig. 2B, contours). As a matter of notation, we refer to MSM-assigned macrostates corresponding to a specific ensemble of conformations with script-text (e.g. *Up*) to distinguish from abstract notions of states, which we quote (e.g. “up”). We investigate the kinetics and thermodynamics among the macrostates (Fig. 2B, arrows). We find a highly-populated *Down* state in exchange with an intermediate *I*_2_, differing by a partial unfolding of the M2-M3 loop (Fig. S2). We find a metastable intermediate *I*_1_ that rapidly relaxes to *I*_2_ (Fig. S3). We find a kinetically-distant *U*_*p*_ state that can transition to and from each of the other three states (Fig. 3A). The TREK-2 system is known to form fenestration sites between the membrane embedded helices wherein inhibitors may bind. The volume of these sites varies monotonically along the up-down coordinate (Fig. SI), with the largest corresponding to the *Down* states.

**FIG. 2.**
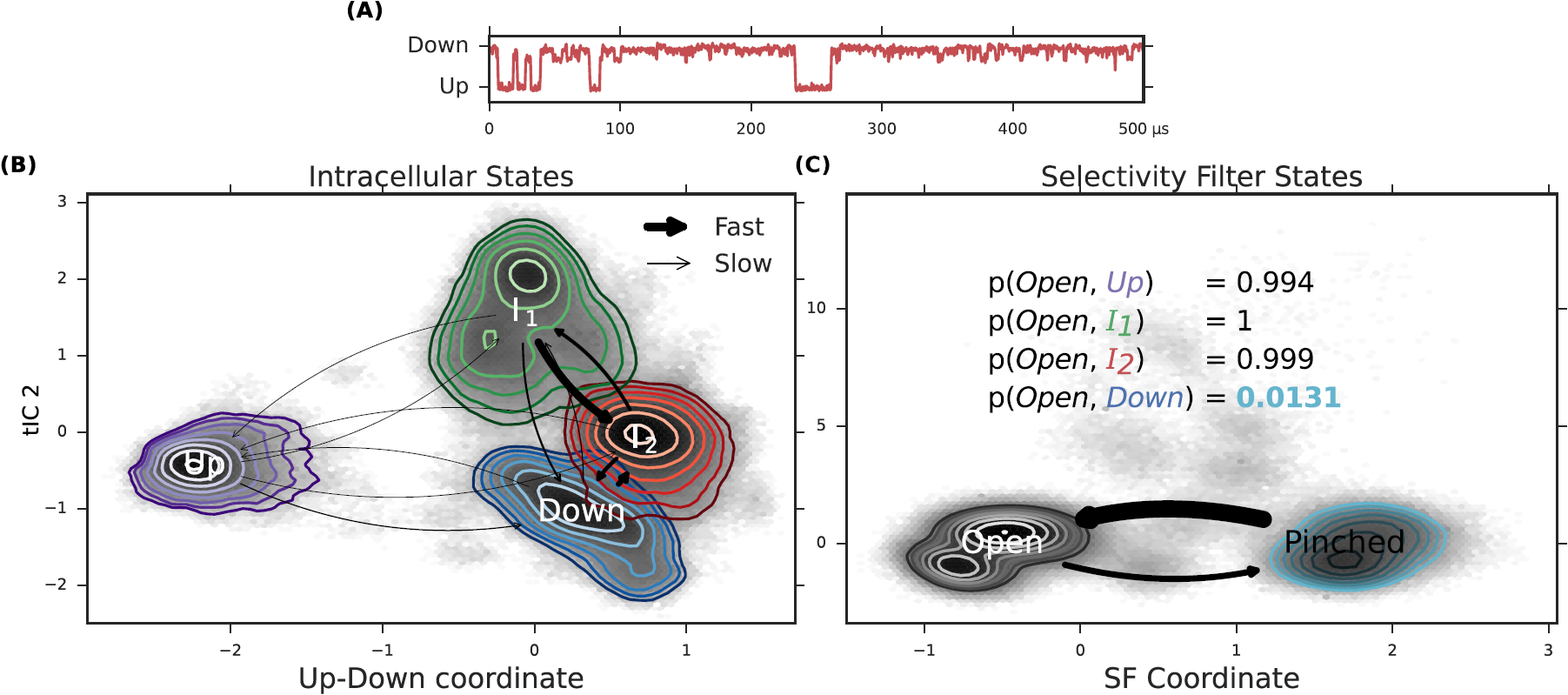
Automatic coordinate discovery using molecular dynamics data identifies an up-down coordinate from a set of highdimensional input data. Modeling the observed dynamics by construction of an MSM permits rapid analysis of conformational dynamics. From our analysis, we discover a novel selectivity filter (SF) conformation. **(A)** MSM analysis permits generation of a 500 μs representative trajectory which contains transitions between up, down, and intermediate conformations, see movie in SI. **(B)** We find four metastable conformational macrostates, shown as contours in tIC space. Transition rates show a rapid relaxation from *I*_1_ to *I*_2_, and a fast exchange between *I*_2_ and *Down*. **(C)** Automatic coordinate discovery on the selectivity filter finds a structurally distinct pinched state. Inset enumerates the SF open probability as a function of intracellular macrostate. The *U*_*p*_ and intermediate intracellular states favor an open SF, while the *Down* intracellular state strongly favors a pinched SF.

**FIG. 3.**
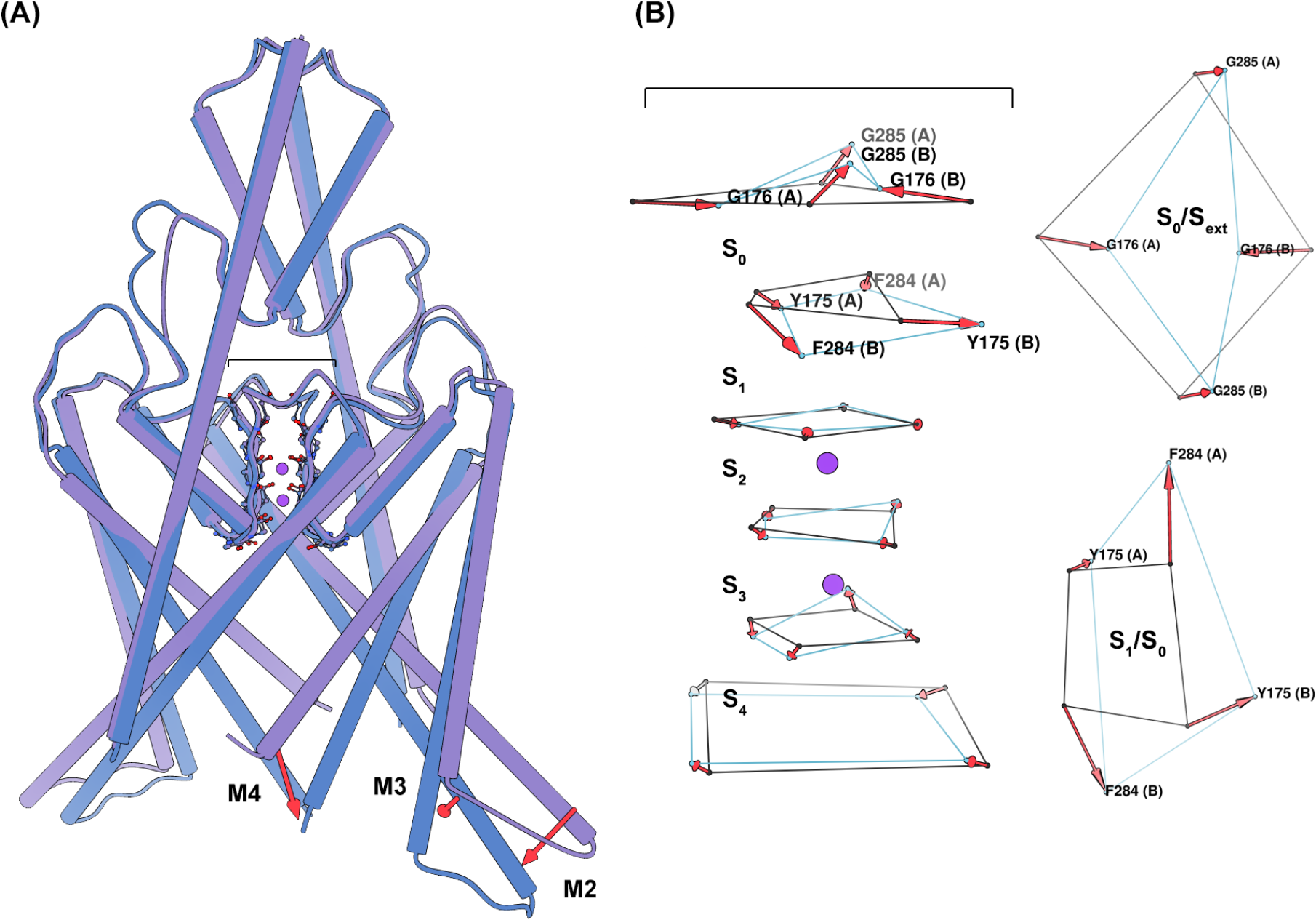
Conformational sampling of the MSM provides structural insight into the major states assumed during simulation. **(A)** A side-view of a superposition of the TREK-2 *U*_*p*_ (purple) and *Down* (blue) intracellular macrostates, highlighting significant conformational differences. Transition from the *U*_*p*_ to the *Down* state involves the large-scale inward movement of M_2_, M_3_, and M_4_. **(B)** A side-view (left) of a superposition of the TREK-2 SF *Open* (black) and *Pinched* (light blue) states, and aerial view (right) of the carbonyl oxygens at the interface of the *S*_1_/*S*_0_ and *S*_0_/*S*_*ext*_ binding sites. Here the SF is represented only by the carbonyl oxygens constituting the SF binding sites. Transition to the *Pinched* filter involves rotations of the residues surrounding the *S*_1_ and *S*_0_ binding sites. These changes disrupt the symmetry of the filter such that the *S*_0_/*S*_1_ carbonyls no longer face the ion conduction pathway (reducing the ability of these sites to coordinate ions), and the extracellular mouth of the filter is reduced in diameter by nearly 4 angstroms (G176 (A) - G176 (B)).

In addition to using tICA to discover an up-down coordinate, we focused the algorithm specifically on the selectivity filter region of the channel. Specifically, we use interatomic distances only among selectivity filter atoms as the input features to learn a second tICA model. In this case, the molecular dynamics reveals a metastable “pinched” conformation of the selectivity filter (Fig. 3B). This SF conformation is not represented in the TREK-2 crystal structures, and could describe a conformation similar to that assumed during C-type inactivation [16]. In this conformation, residues at the top of the *S*_1_ binding site rotate outwards, away from the ion conduction pathway, while residues at the top of the *S*_0_ binding site move inward. This inward movement is most drastic for G176 (A) and G176 (B) (where (A) signifies the protein chain), which restricts the pore from approximately 8.5 to 4.5 angstroms. We model the dynamics between two selectivity filter states (*Open* and *Pinched*) with a Markov model. The model suggests a free-energy minimum *Open* conformation and a metastable *Pinched* conformation (Fig. 2B). Comparing the structural macrostate and SF state for each conformation indicates that the *Pinched* SF conformation is strongly coupled to the channel’s intracellular conformation. The *Pinched* SF state is more prevalent in the fully *Down* state by a ratio of 75:1 (Fig. 2C, inset table). In the *U*_*p*_ and *I*_2_ macrostates, the situation is reversed; the proportion of *Pinched* conformations observed is strongly disfavored and the *Open* configuration dominates. In the *I*_1_ state, only the *Open* SF is seen. The disruption of symmetry and restriction in pore diameter caused by the rearrangements of upper-SF carbonyl oxygens in the *Pinched* state strongly suggests that this conformational difference will negatively affect ion conductance rates. In the following sections, we will provide evidence for this reduced conduction rate.

### B. Dynamics of selectivity filter ion occupancy

TREK-2 not only exhibited large-scale conformational differences in the available crystal structures, but also subtle differences in ion binding. The “down” structures were crystallized with 3K^+^ while the “up” structure was crystallized with 4K^+^. Dong *et al*. [8] hypothesized that all four combinations of (up, down) × (3K^+^, 4K^+^) states are accessible, despite only observing the two. By analogy to other channels, they hypothesized that 4K^+^ states are highly conductive and 3K^+^ states are less conductive.

We test these hypotheses with our MD dataset. Specifically, we investigate the effect of conformational macrostate on ion occupancy in the selectivity filter (SF). By first partitioning the data by conformational macrostate (*Up, Down, I*_1_, and *I*_2_), we construct four MSMs of ion dynamics to compare and contrast the observed ion microstates and rates (shown in Fig. 4A and C). Thick arrows represent rapid transitions. Fig. 4A is the kinetic model for SF ion transitions in the *U*_*p*_ conformation while Fig. 4C is for *Down* (see Fig. S7 and 8 for *I*_1_ and *I*_2_). The relative populations of each ion microstate can be related to the up-down coordinate and is shown as a 2D-histogram in Fig. 4B.

**FIG. 4.**
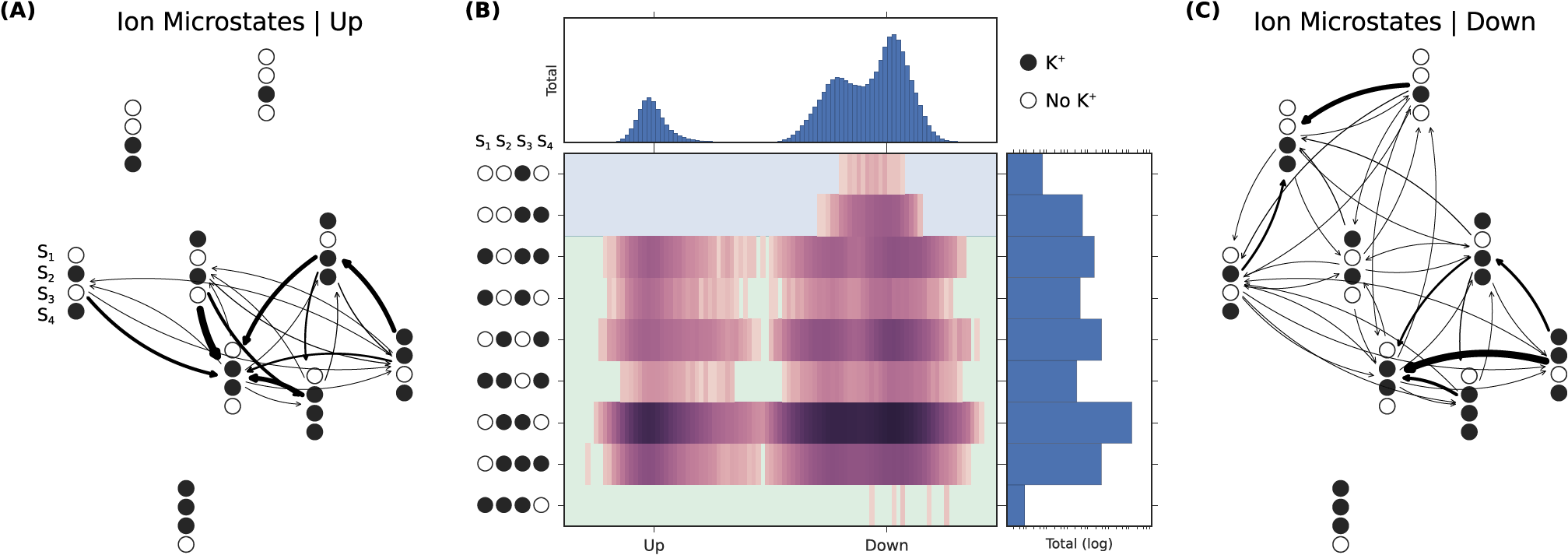
Ion conduction can be modeled as transitions among ion-occupancy states. **(A)** Transition rates between ion states in the *U*_*p*_ macrostate conformation. **(B)** Populations of ion states as a function of the up-down coordinate identified by statistical analysis of the large-scale conformational dynamics. Relative free energies of the ion microstates are conformationally dependent. Additionally, some ion microstates are observed that exist solely in down conformations. **(C)** Transition rates between ion states in the *Down* macrostate conformations.

Several trends are observed in these network graphs and population histogram. For all macrostates, OXXO [X = occupied; O = vacant; positions are given from *S*_1_ to *S*_4_ left-to-right] is found to be the dominant equilibrium ion microstate regardless of macrostate. The OXXO state serves as a sink with many observed transitions into it. This suggests that this state could represent the resting occupancy state. The higher ion occupancies observed in the crystal structures may be an artifact of the cryogenic crystallography conditions (investigated in Ref. [17]. In particular, Fig. S2 of its Supplementary Information). An additional commonality lies in the solvation of the SF ions. The hydration of the ions in the SF during the conduction process is an area of recent contention [18]. The soft knock-on mechanism posits that the ions are never directly adjacent, and they must be padded by at least one water molecule. The hard knock-on mechanism asserts that ions are preferably adjacent to one another. Both theories are based on the knock-on mechanism first formulated by Hodgkin and Keynes in 1955 [19], wherein conduction is explained by the energetic perturbation of an ion entering the SF, and differ only with respect to ion solvation. We find that the full set of probable occupancy states determined from our simulations contains exclusively unsolvated ions. In fact, we observe very little water occupancy for any SF binding site, not just at the sites between ions, for all macroscopic conformations. We find the hard-knock mechanism described by Kopfer *et al*. [17] was supported by our simulations due to the degree of ion solvation and similarity between the set of most probable SF occupancy states. Note that TREK-2 is not canonically voltage sensing [20], so we would not expect the protein conformation to change in the presence of a membrane potential. Hence, we would not expect the populations of the SF occupancy states to change, but we would expect the rates.

The dissimilarities among the macrostate-specific MSMs are striking. The *Down* macrostate displays more diffuse ion dynamics with a large number of possible transitions among an extended state space. The equilibrium OXXO behaves less as a network sink with comparatively fewer high flux transitions into it. One might hypothesize that the greater ion movement would imply lower kinetic barriers and higher conduction. This is counterintuitive as the down state has been suggested to be the low-conductivity state. While it is not clear how or to what extent the large, slow changes in macrostate impact the fast microstate transition pathways and stabilities, this joint conformation-ion analysis highlights qualitative differences that merit further study. To investigate quantitatively the effects of conformation on ion conduction, we perform additional simulation to study specific transitions in more detail.

### C. Impact of structure on function

Large-scale conformational changes among macrostates (Section I A) are suggested to impact the ion transition pathways in the selectivity filter (Section IB), and therefore impact conduction. To quantify these differences, we partition the data by macrostate. For each macrostate, we compute PMF curves for key ion occupancy transitions. The OXXO *→* XXXO and XXOX *→* XOXX transitions were of particular interest because of their proposed role in the hard knock-on ion conduction mechanism.

The OXXO → XXXO transition describes the process of an ion moving from the intracellular channel cavity into the SF. This transition has been proposed to be most critical for the initiation of a conduction cycle. As can be seen in Fig. 5A left, the forward and reverse rates for this process are all less than 3 kCal mol^−1^. The *Up* state is found to have the highest forward kinetic barrier, followed by the two intermediate states, with the *Down* states having the lowest. The reverse rates for each macrostate are comparable, but the sample from the conformation with the alternative SF conformation, *Dowri*_pinch_ displays a highly destabilized second binding well.

**FIG. 5.**
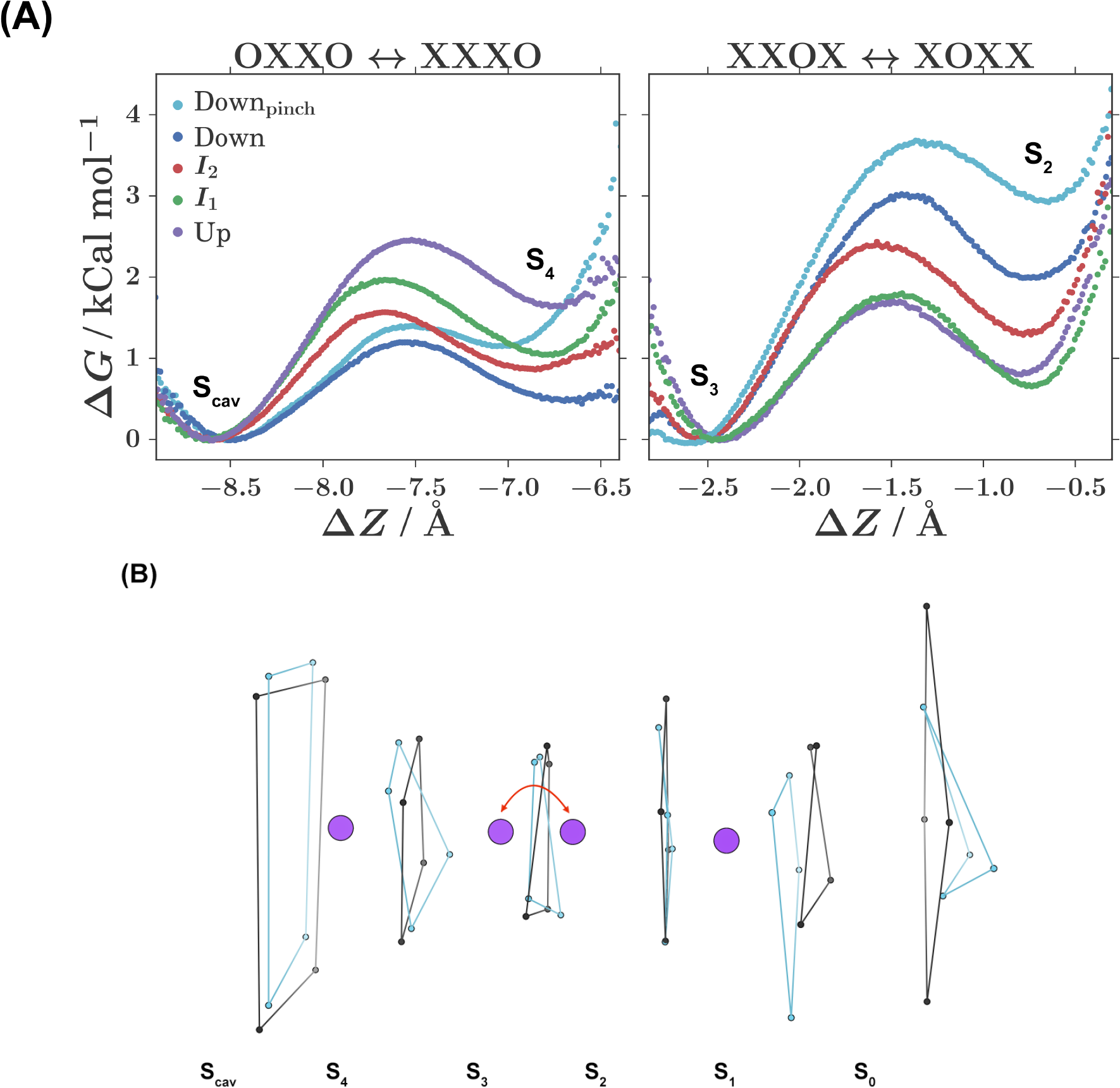
The energetics of ion occupancy transitions are conformationally dependent and can be analyzed using potential of mean force (PMF) calculations. For each conformational macrostate, PMF curves are computed for two key ion occupancy transitions. **(A)** Representative PMF curves for each macrostate. The *Down* macrostate favors the novel, pinched SF configuration and has been separated into *Down* (blue) and Down_p_i_nc_h (light blue) curves. Δ*Ζ* is progress along the ion transition path. **(B)** Illustration of the process sampled during the *XXOX* ↔ *XOXX* PMF calculations for the *Up*_*open*_ (black) and *Down*_*pinched*_ SF states. This process represents the oscillation of an ion between the *S*_3_ and *S*_2_ binding sites, while occupancy of *S*_4_ and *S*_1_ remains static.

The XXOX → XOXX transition describes the movement of an interior ion nearer to the SF exit, from S_3_ to S_2_. This transition was of interest because the conformations sampled from the MSM macrostates displayed variability in the structure of the SF on the extracellular side. This transition is closer to the region of SF structural variability. As shown by Fig. 5A right, the forward and reverse rates for this process are all less than 4 kCal mol^−1^. The size of the forward barrier is highest for the Down states, followed by I_2_, while I_2_ and the up state are the lowest. Once again, the Down_p_¡_nc_h sample yields a highly destabilized second binding well.

Although the *Down* state represents the least conductive state, for the OXXO → XXXO transition it displays the smallest forward kinetic barrier, and for the XXOX → XOXX transition it displays the largest forward kinetic barrier. This illustrates how the conductance rate can be understood only through the consideration of multiple transition events. Barriers are not uniformly higher for the *Down* state. Instead, the effect is stronger for our analyzed downstream conduction transition, especially in the case where an conformational change in the SF has occurred.

For both transitions, conformational macrostate is found to have a marked impact on the forward kinetic barriers, while SF conformation is found to have a marked impact on the reverse kinetic barriers. Alternative SF conformations are observed with the highest probability for the *Down* state from the MD dataset (see Fig. 2C and SI). The conformational changes associated with these alternative SF states are characterized by rotations of the backbone dihedral angles for the residues at the interface of the *S*_1_ and *S*_0_ binding sites (see Fig. 3B). In the case that a backbone dihedral rotation did occur, the *S*_1_ carbonyl oxygens no longer face the ion conduction pathway. Because the carbonyl oxygens of the residue backbone are responsible for coordinating the ions at each binding site, these rotations could render *S*_1_ nonfunctional. Additionally, these rotations allow for the inward movement of the *S*_0_ G176 (A) and G176(B) residues. This movement drastically restricts the diameter of the SF extracellular mouth, which could present a steric barrier to ion permeation.

As evidenced by the PMF curves, the *Pinched* SF conformations disfavor the extracellular movement of ions through the SF, and could represent a state similar to SF inactivation. The p-loop strand constituting the side of the SF with the most frequent rotation (F284B) is tethered to M4. This helix region displays a stark conformational difference between the *U*_*p*_ and *Down* states, and terminates at the c-terminal regulatory domain. We propose that the movement of M4 as the protein assumes the *Down* state allows for a conformational change of the SF *S*_1_ ion binding site, resulting in disfavored conduction. Coupling between c-terminal regulation and SF gating has been demonstrated for Kir [21] and KcsA [22] potassium channels, and has been suggested to apply to K2P channels as well [6, 23]. Our results provide further evidence for this claim.

## II. DISCUSSION

We applied molecular dynamics and MSM analysis to predict that a novel pinched state favored during membrane compression represents a low-conducting state of the TREK-2 K2P potassium channel. An initial hypothesis [8] suggested a direct connection between intracellular motion and channel activity. Membrane stretch and compression (“up” versus “down” macrostates) would beget high- and low-conducting functionality, respectively. This hypothesis has been suggested to be overly simplistic by experimental measurements [4] and the present study. After simulating the TREK-2 channel for 195 μs, we discover a novel selectivity filter conformation highly preferred by the *Down* state, but disfavored by the down-like *I*_1_ and *I*_2_ metastable intermediates and *Up* state. Steric consequences of this SF configuration suggest an impact on conductance. For further investigation, we use MSMs to model the behavior of ions in the selectivity filter conditioned on the conformational macrostate of the channel. Our ion dynamics models highlight similarities between the kinetic networks from the different conformations.In particular, the Kopfer *et al*. [17] hard-knock mechanism is well supported by our simulations. The macrostate-conditioned models also suggest striking qualitative differences between ion microstate occupancies and transition paths. To gain a quantitative understanding of these differences, we selected representative conformations from our unbiased MD calculations and computed PMF free energy profiles for crucial ion transitions, again partitioned by conformation. While the initial transition OXXO → XXXO shows the smallest barrier for down configurations (contrary to the idea that down conformations are less conductive), a later Kopfer pathway transition shows the highest barrier for down states, particularly the down state with pinched selectivity filter (*Down*_pinch_). The *Down*_pinch_ state is shown to have destabilized second wells in both important transitions studied, adding further evidence to the reduced activity of this conformation. Our findings agree with the growing consensus that intracellular motion is not directly coupled to conduction. Instead, conformational states affect selectivity filter conformations. We contribute a structural basis for this idea by the discovery of a novel pinched SF state involving rotations around Y175/F284. We encourage future structural studies to target this novel state.

## III. METHODS

### A. Molecular Dynamics

The simulations were started from four crystal structures of TREK-2 (PDB codes: 4xdj, 4bw5, 4xdl, 4xdk) [8] Crystal structures were solvated in a 4:1 POPC:POPE lipid bilayer membrane using charmm-gui [24]. Simulations inputs were generated using the tip3p [25], lipid14 [26] and AMBER14SB [27] forcefields. Equilibration was performed with restrained proteins and lipids, by slowly increasing temperature in amber [28]. Production simulations were run on Folding@Home [10] using OpenMM 6.2 [29] and Gromacs 4.5.3 [30]. PMF calculations were run with Gromacs 5.0.5 on Bluewaters and reweighted with WHAM [31]. Analysis was performed with MDTraj 1.5 [32] and MSMBuilder 3.4 [33]. Conformations were visualized with VMD [34] and Chimera [35].

### B. Markov State Modeling

The pore region of the protein was featurized by taking all respective chain A to chain B distances between residues 1-24, 112-189, 230-256. These features were transformed into kinetic coordinates with tICA (lag-time = 1 ns). Conformations were clustered into 500 states using the first 3 dimensions of tIC coordinates via the Mini-batch KMeans algorithm. An MSM was fit using MLE at a lag-time of 20 ns. Conformational microstates were lumped into four macrostates with PCCA+. Ion occupancies were featurized by enumerating the 16 possible occupancy states (from four binding sites). An MSM was fit using MLE, again at a lag-time of 20 ns.

### C. PMF Calculations

PMF initial conformations were obtained through PCA analysis on the protein dihedral angles for each MSM state. These conformations were aligned such that the SF was centered about the z-axis. Pulling simulations were performed in order to sampling the reaction coordinate for each ion transition. Five umbrella sampling simulations were run for each PMF, one in each ion binding well, one at the transition state, and one between each binding well and the transition state. These simulations were run with a spring constant of 8000 kCal mol^−1^ for 6 ns, with the first 1 ns treated as equilibration. For each transition and macrostate, at least 8 conformations were analyzed. The PMF reaction coordinate was defined as the projection of the distance from the SF center of mass to the transitioning ion onto the SF symmetric axis. All PMF related simulations were run using Gromacs 5.0.5 [30]. RMSD of protein conformation to initial structure was monitored during umbrella sampling simulations to ensure there were no artifacts caused by the umbrella potential.

## IV. ACKNOWLEDGMENTS

We thank John Mathias and Mark Bunnage for their encouragement and support. We thank Arianna Peck and Brooke Husic for critical feedback on the manuscript. We acknowledge funding from NIH grants U19 AI109662 and 2R01GM062868. This work is funded by Pfizer’s Science and Technology budget. We thank the Folding at Home donors who contributed to this project (PROJ9712, 9761 and 9762). This work is part of the “Petascale Simulations of Biomolecular Function and Conformational Change” PRAC allocation support by the NSF (award number 1439982).

## V. AUTHOR CONTRIBUTIONS

All authors designed the research. MPH and KAM performed the research. RAD and VSP supervised the research. All authors edited and approved the manuscript.

